# A segregating inversion generates fitness variation in a yellow monkeyflower (*Mimulus guttatus*) population

**DOI:** 10.1101/028670

**Authors:** Young Wha Lee, Lila Fishman, John K. Kelly, John H. Willis

## Abstract

Polymorphic chromosomal rearrangements, which can bind together hundreds of genes into single genetic loci with diverse effects, are increasingly associated with local adaptation and speciation. They may also be an important component of genetic variation within populations. We genetically and phenotypically characterized a novel segregating inversion (*inv6*) in the Iron Mountain (IM) population of *Mimulus guttatus* (yellow monkeyflower). We first identified a region of recombination suppression in three F_2_ mapping populations resulting from crosses among IM plants; in each case, the F_1_ hybrid parent was heterozygous for a homogenous derived haplotype (*inv6*) across markers spanning over 4.2 Mb of Linkage Group 6. Genotype-phenotype associations in the three F_2_ populations demonstrated negative *inv6* effects on male and female fitness components. In addition, *inv6* carriers suffered a ~30% loss of pollen viability in the field. Despite these costs, *inv6* exists at moderate and apparently stable frequency (~7%) in the natural population, suggesting counter-balancing fitness benefits that maintain the polymorphism. Across four years of monitoring in the field, *inv6* had an overall significant positive effect on the seed production (lifetime female fitness) of carriers. This benefit was particularly strong in harsh years and may be mediated (in part) by strong positive *inv6* effects on flower production. These data suggest that opposing fitness effects maintain an intermediate frequency, and as a consequence, *inv6* generates inbreeding depression and high genetic variance. We discuss these findings in the context of theory about the genetic basis of inbreeding depression and the role for chromosomal rearrangements in population divergence with gene flow.

## Summary

Polymorphic chromosomal rearrangements may contribute importantly to genetic variation within populations. Here, we identify a novel segregating inversion (*inv6*) in that contributes to fitness variation in a natural population of *Mimulus guttatus* (yellow monkeyflower). Greenhouse studies demonstrate negative *inv6* effects on male and female fitness components. Despite these costs, *inv6* exists at moderate and apparently stable frequency (~8%) in nature, and four years of field data, indicate significant positive effects on female fecundity. Opposing fitness effects apparently maintain an intermediate frequency, and as a consequence, *inv6* generates substantial genetic variance in fitness.

## Introduction

Polymorphic chromosomal inversions are an important component of genetic variability (Hoffmann and Rieseberg 2008). Inversion polymorphisms are associated with species differentiation in both flowering plants (Rieseberg et al. 1999; Fishman et al. 2013; Hermann et al. 2013) and animals (Noor et al. 2001). Within species, inversion polymorphisms often exhibit clinal variation suggestive of adaptation to latitudinal environmental variables (Balanyà et al. 2003; Hoffmann and Rieseberg 2008; Fang et al. 2012; Cheng et al. 2012). Similarly, putatively adaptive differences in traits related to fitness among populations or ecotypes co-segregate with inversions within many species, including Drosophila (Krimbas and Powell 1992), Anopheles mosquitoes (Coluzzi et al. 2002), Rhagoletis flies (Feder et al. 2003), seaweed flies (Gilburn and Day 1999), monkeyflowers (Lowry and Willis 2010), and sticklebacks (Jones et al. 2012). These patterns support the idea that inversions contribute to local adaptation and speciation because they suppress recombination among multiple genetic variants with context-dependent effects on fitness (Kirkpatrick and Barton 2006).

The same evolutionary processes that generate differences among geographical populations can also maintain chromosomal polymorphisms within populations. Environmental fluctuations or frequency-dependence could allow the persistence of alternative arrangements containing sets of co-adapted alleles, each set being optimal under one or another condition (Dobzhansky 1970). This hypothesis may account for the “supergenes” that segregate within some species, such as those underlying mating types in heterostylous plants or wing mimicry polymorphisms in *Heliconius* butterflies (reviewed in Schwander et al. (2014) and Thompson and Jiggins (2014)). However, a simpler alternative scenario for polymorphism is that a newly arising chromosomal rearrangement may happen to contain one or more intrinsically detrimental alleles along with an advantageous allele (Kirkpatrick & Barton, 2006). Theory suggests that recessive or partially recessive deleterious alleles should be individually rare, but they are common as a class of variant (Muller 1918; Sturtevant and Mather 1938; (Kondrashov 1988) and thought to be the major cause of inbreeding depression (Charlesworth and Willis 2009). If the beneficial variants caught by a new inversion increase fitness as heterozygotes (i.e., are at least partially dominant), whereas the detrimental alleles are at least partially recessive, a stable polymorphism may result. The novel inversion will increase in frequency initially, as its deleterious recessive alleles will be rare and thus hidden in heterokaryotypes. However, as the inversion frequency rises, recessive costs are expressed more often. Eventually, the benefits and costs balance, and an equilibrium frequency will be maintained. Thus, rather than promoting adaptation, the inversion brakes the spread of a beneficial allele and leads to elevated frequencies of deleterious variation.

Selfish genetic elements, which are routinely polymorphic with species, illustrate how suppression of recombination among loci with opposing fitness effects can lead to balanced polymorphism (Burt and Trivers 2008). In these cases, the positive selection comes from a transmission advantage rather than a benefit to heterozygote fitness, but the effect is similar. A striking instance of such a balanced polymorphism is the female meiotic drive locus of *Mimulus guttatus* (common yellow monkeyflower) (Fishman and Saunders 2008; Fishman and Kelly 2015). A structural variant of the centromeric region of chromosome 11, known as *D*, exhibits centromere-associated drive over the alternative chromosomal type (*D*^−^) in the Iron Mountain, OR study population. D gains a ~60:40 transmission advantage in heterozygote individuals by driving through female meiosis. Consistent with this selective advantage, patterns of nucleotide diversity suggest a recent and rapid spread of the *D* variant at Iron Mountain. However, *D* is prevented from reaching fixation because it exhibits recessive negative effects on both male (pollen viability; Fishman and Saunders 2008) and female (seed set; Fishman and Kelly 2015) fitness components in nature. Together, these linked recessive costs maintain the *D* chromosomal variant at intermediate frequency (30-40%), near the predicted equilibrium. Because it elevates the frequency of deleterious recessive variants, *D* contributes substantially to both inbreeding depression and the genetic variance in fitness (Scoville et al. 2009).

In this paper, we characterize a second major structural variant putatively maintained by balancing selection within the same Iron Mountain *M. guttatus* population *First,* in each of three independent QTL mapping crosses between six plants sampled from a single population, we discovered a region of extreme recombination suppression, spanning over 4 Mb on the upper end of Linkage Group 6 (LG6). In each cross, we found that the F_1_ hybrid parent was heterozygous for a unique derived haplotype in this region, which appears to correspond to a newly arisen chromosomal inversion (henceforth *inv6*). *Second,* we found that major QTLs for pollen viability and number (along with other fitness components) in each of the three F_2_ populations map to this chromosomal region, indicating that *inv6* has substantial negative effects on male and female fertility under greenhouse conditions. *Third,* we tested whether the *inv6* haplotype is associated with reduced pollen viability in the wild population, and find that it is strongly deleterious for male fertility despite its moderate (7-15%) frequency in wild and wild-derived population samples. *Fourth,* to test the hypothesis that positive associations with other fitness components maintain the *inv6* polymorphism, we examine the effects of *inv6* genotype on female fitness components over four years in the field. We find an overall positive (and likely dominant) effect of *inv6* on lifetime female fitness, mediated in part by strong positive effects on flower production in both years for which we have flower number data. Together, these data suggest that opposing male and female fitness effects the *inv6* chromosomal inversion contribute to the maintenance this structural polymorphism. We argue that inversions under such balancing selection may be common and major contributors to inbreeding depression in natural populations.

### MATERIALS AND METHODS

*Study system* – *Mimulus guttatus* (2n=28, Phrymaceae) is a self-compatible wildflower that grows throughout Western North America. It is the most common member of an eponymous species complex comprised of highly polytypic, partially inter-fertile subspecies (Vickery 1978; Wu et al. 2007). We focus on a population located on Iron Mountain (IM) in the Western Cascades of Oregon. The site is an alpine xeric meadow composed of a steep north facing slope at an elevation of 1470 m, over an area of ~600m^2^. The population usually comprises hundreds of thousands of flowering individuals each year, is bee-pollinated, and has a mixed mating system with an estimated selfing rate of 0-25% (Willis 1993; Sweigart et al. 1999). The population shows no evidence of spatial genetic structure (Sweigart et al. 1999) or biparental inbreeding (Kelly and Willis 2002).

***Mapping populations*** – As part of another study to investigate the genetic basis of complex trait variation and inbreeding depression within the IM population, we crossed 6 plants to produce 3 mapping populations. We conducted two different sets of genetic mapping experiments. The first (the replicated F_2_) involved genotyping markers across the genome, while the second (IM179xIM767) was genotyped only for markers on Linkage Group (LG) 6. The grandparents of the replicated F_2_ were sampled from a selection experiment on flower size (Kelly 2008; Kelly et al. 2013). Selection was sustained for six generations (bi-directional on corolla width) within populations that were maintained at large size. After a generation of random mating without selection within each population, three Low parents were selected from the Low population and each was randomly paired to distinct High parent. We crossed the plants within each High-Low pair and randomly selected a single F_1_ offspring. Three mapping populations, each consisting of 378-384 F_2_ individuals, were derived from selfing the three F_1_ progenitors. These are called the c2, c3, and c4 mapping populations and each F_2_ was scored as HH, HL, or LL at each locus according to the grandparental origin of a marker allele. Following discovery of the inversion on chromosome 6 (see below), we generated an additional cross between two inbred lines from Iron Mountain (IM179 and IM767) with the same orientation of markers on chromosome 6. We selfed a single F_1_ from this cross to produce 86 F_2_ plants that we grew and then genotyped at 19 marker loci.

***Genotypes and phenotypes in the replicated F_2_ experiment*** *–* As part of our original QTL mapping design, we measured days to flower, pollen viability, pollen number, and supplemented seed set on each F_2_ individual in the University of Kansas greenhouses in the Spring of 2003. The on each plant the first and second flower were sampled for pollen while the third and fourth flower were hand-pollinated (with later harvest and counting of seed) to estimate supplemented seedset. We used a common pollen donor (IM62, a standard inbred line derived from Iron Mountain) for all flowers. Pollen number and viability was measured using a Coulter Counter following the protocol of (Kelly et al. 2002). We collected bud meristem tissue from each of 378, 384, and 384 individuals of mapping population c2, c3, and c4, respectively, and extracted DNA using a CTAB procedure (Kelly and Willis 1998).

Nearly all of the markers used for this study (prefixed with MgSTS – *M. g*uttatus *s*equence *t*agged *s*ite – or simply “e” – for *e*xpressed sequence tag) are exon-primed markers spanning introns. To identify informative markers in each cross, the High and Low outbred parents and progenitor individuals were screened for 748 MgSTS markers that had been successfully amplified in IM62, a standard inbred line derived from Iron Mountain. The forward primer was tagged in the 5’ end with fluorescent dye and the resulting labeled PCR products were run on ABI 3730 or 3700 Genetic Analyzers (Applied Biosystems, Foster City, CA). PCR amplification followed a standard touchdown protocol (e.g. see Fishman and Willis 2005) with multiplexing and pooling based on expected allele sizes. We scored genotypes using Genemarker software (Softgenetics, State College, PA) based on the segregation of length-variable alleles. These markers are usually co-dominant and single copy allowing genetics maps from different experiments to be tied together through markers in orthologous genes (MgSTS markers are available from www.mimulusevolution.org). We also genotyped several microsatellite markers (prefixed by “aat”, (Fishman et al. 2001)) and custom-designed markers (prefixed by “yw”) in each mapping population to fill large gaps in the maps (>30 cM).

***Linkage map construction*** – We built the linkage maps following three iterative rounds of data quality control. We used JOINMAP 4.0 (Stam 1993) to construct preliminary maps by regression mapping and examined each marker for amount of missing data and for incidence of double crossovers. Markers with significant missing data (20% or more) were re-amplified depending on whether the marker was critical to fill a gap, otherwise it was discarded. Markers that had significant numbers of double crossovers were either discarded, or had certain problematic individuals coded as “missing”, or was re-coded as dominant, or was re-amplified depending on the importance of the marker for map resolution. Special effort was made to identify and place markers that filled in large gaps and markers that were shared between crosses. The final missing data proportions were 4.7%, 2.8 %, and 3.3% for c2, c3, and c4, respectively.

We finalized linkage maps for each mapping population with the Kosambi mapping function using the maximum likelihood algorithm in JOINMAP 4.0 run with default settings. We made two versions of linkage group 6 for each map as this chromosome exhibited extensive recombination suppression involving 20-30 markers in each map. One version was made using the maximum likelihood algorithm as above, with all but one representative marker in the suppressed recombination region deleted. Deleting excess markers in the region of suppressed recombination was necessary for permutation tests of significance. The second version of LG6 was made to illustrate recombination suppression. We included all genotyped markers on LG6 and performed regression mapping. Taking the ends of linkage groups into account, total map length was calculated by adding 2s to the estimated length of each linkage group, where s equals the average inter-marker distance for that linkage group. Assuming random distribution of markers, estimated genome coverage was calculated as c=1-(e-2dn/L), the proportion of the genome within distance d of a marker, where n is the number of markers and L is total map length.

***Quantitative Trait Locus (QTL) mapping and inv6 phenotypic effects*** *–* For genome-wide mapping of fertility traits in the three F_2_ populations, we used composite interval mapping in Windows QTL Cartographer 2.5 (http://statgen.ncsu.edu/qtlcart/WQTLCart.htm) to set priors for Bayesian mapping as implemented in Rqtlbim (Yandell et al. 2007; Yi et al. 2007). Details of the QTL mapping methods are given in the Supplemental Methods. To estimate the effects of inv6 on fertility traits in the mapping populations, we used ANOVA with *inv6* genotype inferred from diagnostic markers with ambiguous genotypes were treated as missing data.

***Identification of the inversion haplotype*** *–* We grew and extracted DNA from a sample of 96 outbred individuals from the Zia-1 base population, the source population for the artificial selection experiment (Kelly 2008). We genotyped each sample for 18 markers spanning LG6, both in recombination-suppressed and freely recombining regions.

***Field measures of male and female fitness components*** *–* In 2007, we collected the pollen from all four anthers of newly opened flowers (1 per plant) in the Iron Mountain *M. guttatus* population, and then harvested plants for later DNA extraction (Fishman & Saunders 2008). Pollen was stained with aniline blue, and a subset of fertile and sterile grains counted using a haemocytometer. Individuals (n =187) were genotyped at e423 and e723, markers diagnostic for inv6. Genotypic effects on male fertility traits were analyzed with t-tests in JMP. In 2010-2014, we collected entire senescing *M. guttatus* plants at the Iron Mountain population, then recovered and counted their fruits, seeds, and (in 2012 and 2013) flowers (Fishman and Kelly 2015). We then extracted genomic DNA from the remaining tissue, and genotyped individuals at the inv6-diagnostic marker e423 (n = 1248 over the 4 years). Because we only collected plants that survived to mature at least one fruit, our measures of female fitness do not include survival-to-reproduction.

Because there were too few *inv6* homozygotes per year to include all three genotypes in the analyses, we coded individuals as *inv6* carriers if they were either heterozygotes (one inv6-diagnostic 236bp allele at e423) or *inv6* homozygotes. This allows us to evaluate dominant and/or additive effects of the inversion, but not recessive ones. We examined female reproductive trait variation (flowers, fruits, seeds/fruit, and total seeds) using generalized linear models in JMP 11 (SAS Institute, Cary NC), with year and *inv6* as a main effects. For the first three traits, we used an over-dispersed Poisson distribution (loglink function), whereas we analyzed log-transformed seed number (+1) using a normal distribution and identity link (as in Fishman and Kelly 2015).

The plants genotyped for *inv6* only partially overlapped those genotyped at the *D* female meiotic drive locus, which affected several female fitness traits over the same seed collections (Fishman and Kelly 2015). To maximize sample size per year, we did not include *D* genotype in these analyses; however, we verified that there was no statistical association between the two polymorphisms (Pearson χ2, *P* = 0.61). There were never significant year x genotype interactions (lowest *P* = 0.09 for log[total seeds]), so we present analyses without interaction effects. However, our study period spanned two fairly standard growth years (2010 and 2012: 1-2 fruits, 40-60 seeds per plant) (as in Fishman and Willis 2008), and two relatively lush years (2011 and 2013). Using mean seedset as a proxy for length of the growing season, we then whether the inversion had different effects in good (mean seedset 100-250) and poor (mean seedset 40-55) years, in an analysis with quality, year (quality), *inv6*, and *inv6* x quality interaction.

## RESULTS

***Discovery of* inv6 *region by recombination suppression—***All three of the replicate High x Low F_2_ genetic maps (Supplemental Figure S1) revealed a large cluster of markers on Linkage Group 6 (LG6) that appeared to be completely linked. This non-recombining region, spanning marker e25 to e804 on the consensus map (Figure 1), corresponds to ~4.2 Mb of the v2.0 *M. guttatus* physical map (http://phytozome.jgi.doe.gov/). In contrast, the same set of markers corresponds to 40 cM in the IM79xIM767 cross (Figure 1), as well as in other *M. guttatus* complex linkage maps (Lowry and Willis 2010; Fishman et al. 2014). Each High x Low F_2_ segregated for a particular shared haplotype (i.e. a specific set of alleles) at contiguous MgSTS loci across the recombination-suppressed region. This haplotype, which we denote as *inv6,* derived from the Low grandparent (small flower size) in c2 and from the High grandparents in c3 and c4, suggesting that its presence in all three crosses was not related to divergent floral selection.

**Figure 1.**
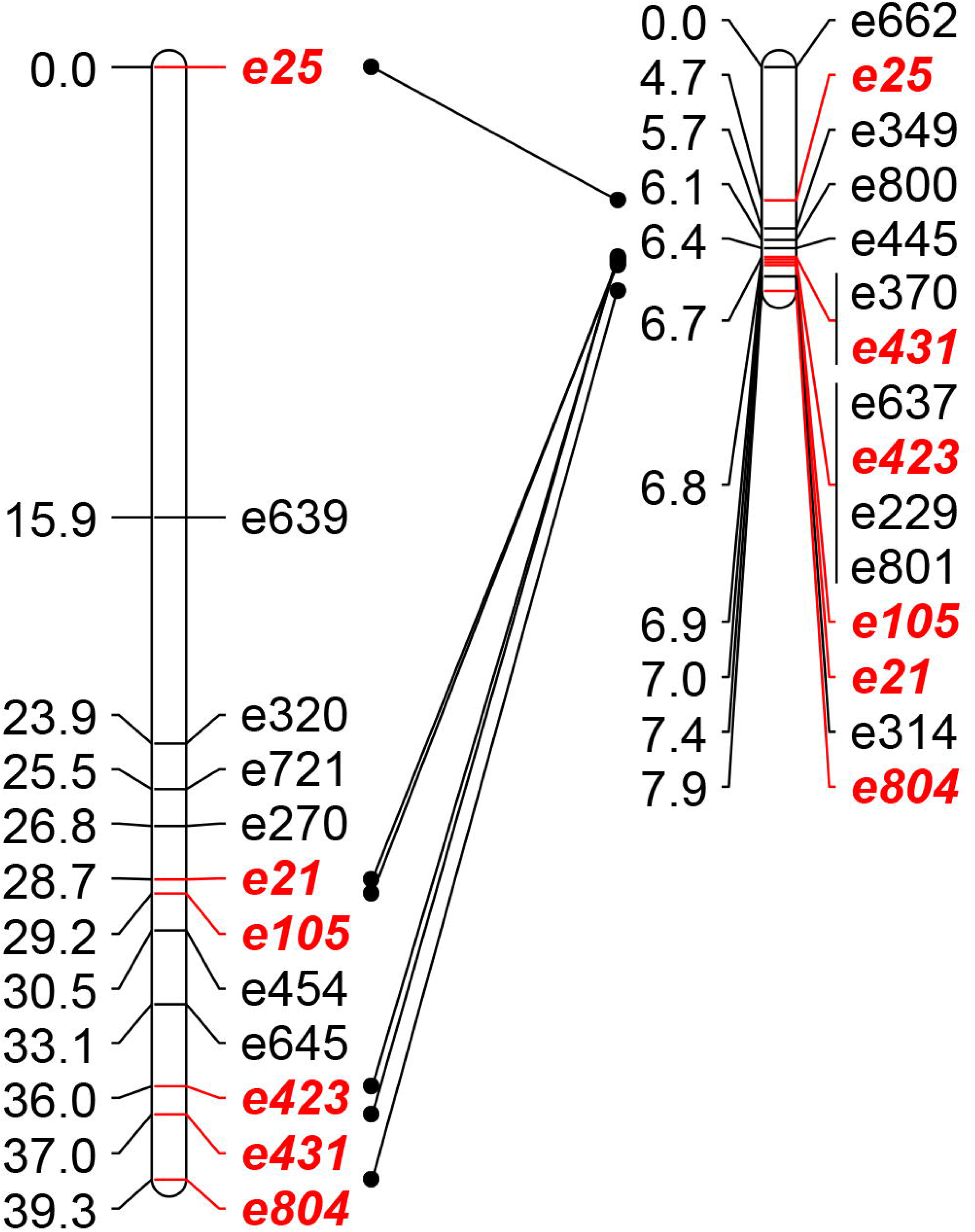
Comparative linkage mapping of the upper end of LG6 in Iron Mountain *Mimulus guttatus* hybrids. In a freely recombining cross (left: IM179xIM767 F_2_; N = 96), this region spans ~40cM, whereas recombination is highly suppressed in all three F_2_ mapping populations segregating for the *inv6* haplotype (right; c3 map shown). Marker names to right, cM to left of bar. Shared markers are highlighted (red, italic).

***QTLs for fitness traits map to Inv6***– *Inv6* affected trait means differently in each of the three mapping populations (presumably due to variation in the non-*inv6* alleles and the different genetic backgrounds), but all significant effects were negative (Figure 2). In c2, the inversion reduced total pollen (F_2, 369_ = 11.26, *P* < 0.001), pollen viability (proportion of grains viable; F_2_, 369 = 40.32, *P* < 0.001) and supplemented seedset (female reproductive capacity; F_2, 364_ = 14.87, *P* < 0.001) per flower. For all three measurements, gene action was recessive, with the negative effect limited to *inv6* homozygotes (Figure 2). In c3, there were no effects on female fertility, but a significant negative effect on pollen viability (F_2, 379_ = 9.43, *P* < 0.001). Gene action is more nearly additive in c3 (Figure 2). In c4, *Inv6* had strongly deleterious effects on female fertility (F_2, 343_ = 10.40, *P* < 0.001), total pollen (F_2, 345_ = 10.04, *P* < 0.001), and pollen viability (F_2, 345_ = 92.67, *P* < 0.001). Heterozygotes were intermediate for all three measurements in c4; *Inv6* partially recessive its effects on female fertility and total pollen but slightly dominant for pollen viability.

**Figure 2.**
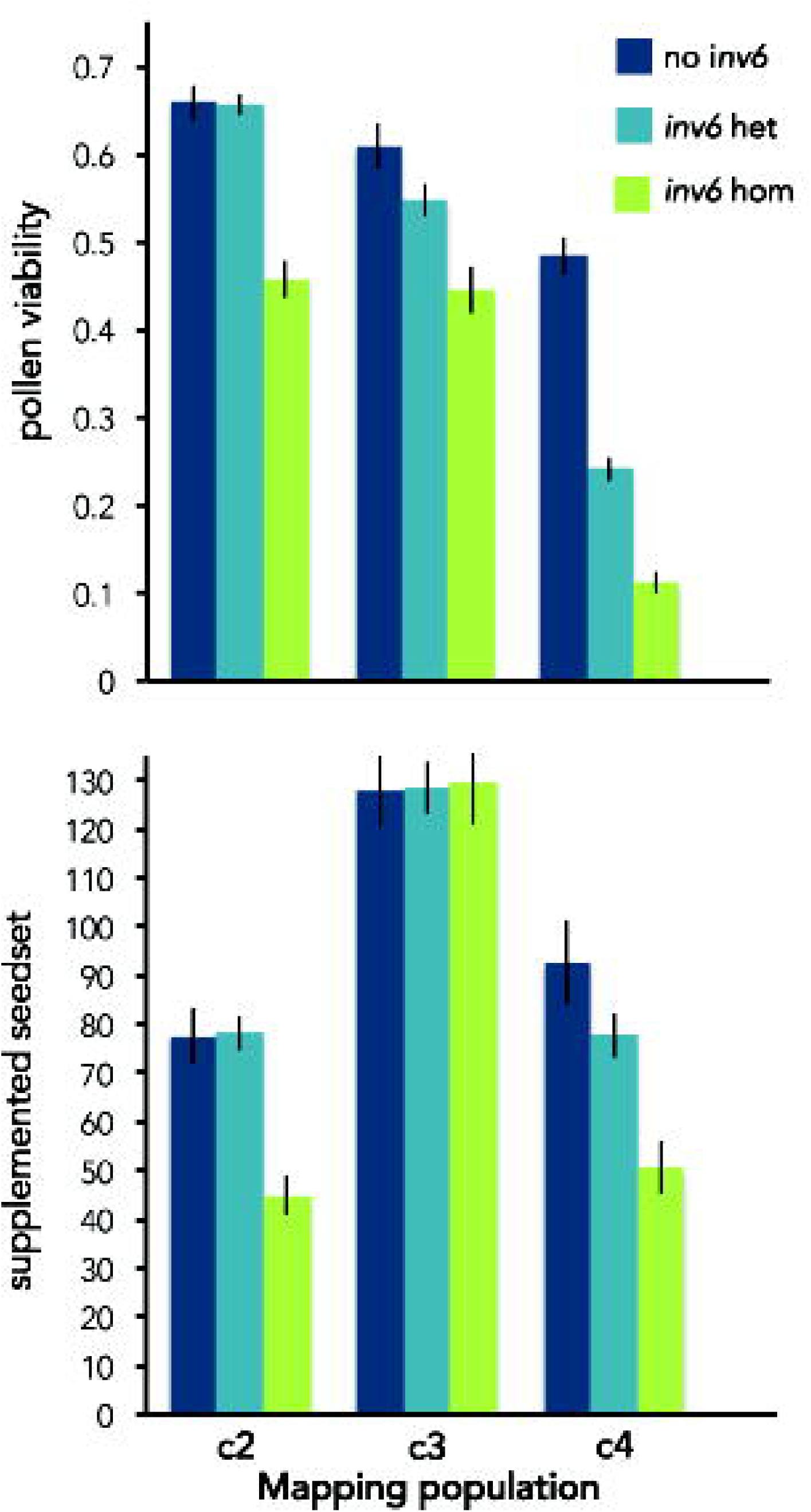
Effect of *inv6* genotype on pollen viability (mean +/− 1 SE) and supplemented seed set within each of the F2 mapping populations.

Overall, trait means differed substantially among the three mapping populations (Supplemental Table S1). Consistent with a highly polygenic basis for fertility variation, we mapped 30 QTLs for pollen number or pollen viability in other parts of the genome, as well as two QTLs for supplemented seed set (Supplemental Table S2). The LG11 meiotic driver *D* also segregated in each cross and, as expected, the drive allele reduced male fitness. Most fertility QTLs exhibit intermediate dominance, but we did map one under-dominant QTL (for viable pollen number) and three over-dominant QTLs (two for viable pollen number and one for supplemented seedset). However, none of these were present in more than one mapping population. Excluding the two major QTLs (*inv6* and the meiotic driver) as well as the over/under-dominant QTLs, low alleles were moderate in effect (average s = 0.31) and partially recessive (average h = 0.18). These estimates, combined with the fact that these QTLs were always segregating in only one of the three crosses, suggests they may be deleterious alleles segregating at low frequencies in the natural population.

***Frequency of inv6 in nature—***In the Zia-1 base population, which was derived via two generations of hand-outcrossing from a wild IM sample of over 1000 plants, the *inv6* haplotype had an overall frequency of 15% (Figure 3). Allelic variation in this population indicates that *inv6* is a derived haplotype, as it shares alleles with the alternative arrangement at many loci. However, in this sample, *inv6* has private alleles at a few apparently diagnostic loci (e.g. e723 and e423). In the flowering plants sampled from the field, the estimated frequency of *inv6* was slightly lower (7-8%, see next section).

**Figure 3.**
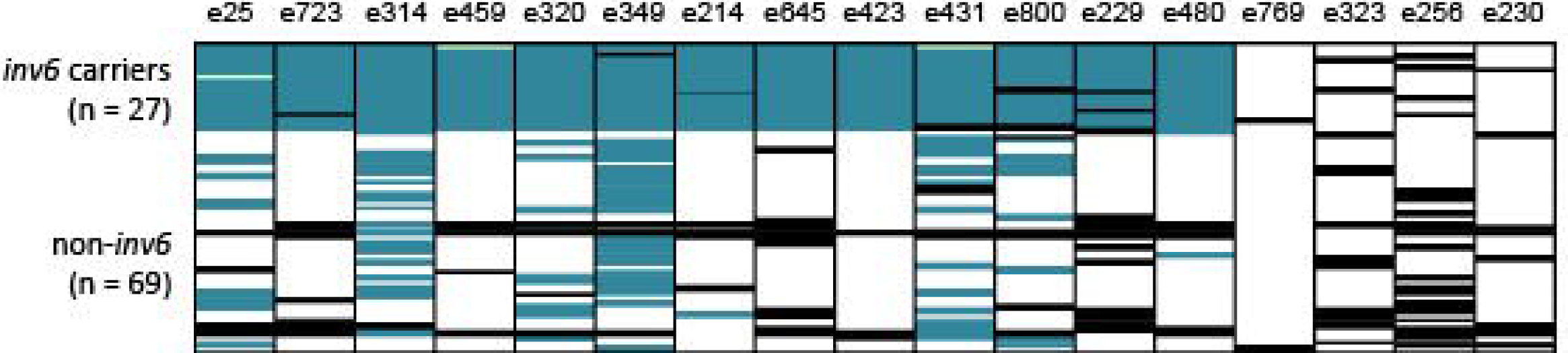
Delineation of *inv6* haplotype block in *M. guttatus* individuals derived from wild IM plants by one generation of outbreeding (n = 96). Each cell represents the genotype of an individual genotyped at 17 markers across LG6. Genotypes carrying at least one *inv6*-associated allele are shown in blue (for the 13 *inv6*-spanning markers) and non-*inv6* genotypes are shown in white. Black = missing data, and three non-*inv6* genotypes (likely genotyping errors or double crossovers) within the *inv6* block are shown in green.

***Inv6 fitness effects in the field –*** Given strong negative effects on fertility traits in the greenhouse mapping populations, the non-trivial frequency of *inv6* in the IM population is surprising. To evaluate both the frequency and the fitness effects of *inv6* more directly, we capitalized on two existing samples of wild IM plants. First, we examined pollen viability and pollen number in field samples collected in 2007. As in the greenhouse experiments (particularly c4, where it was additively deleterious), carriers of *inv6* had 30% lower pollen viability than non-carriers (*P* < 0.0001, N = 187; Figure 4). Homozygotes are too rare in the field to estimate dominance effects. These deleterious effects in heterozygotes make the observed frequency of *inv6* (8% in the 2007 sample of wild plants) extremely unlikely without counterbalancing benefits. Second, we examined effects of *inv6* genotype on female fertility traits (flower number, fruit number, seeds/fruit, and total seed number) from 2010 through 2013. There was strong year-to-year variation (all *P* < 0.0001) in all these traits, indicating that this sample captures the breadth of natural environmental variation in female fertility. Similar to 2007, the frequency of *inv6* was ~7% (~14% heterozygous plants), and was fairly constant across years in the 2010–2013 samples.

**Figure 4.**
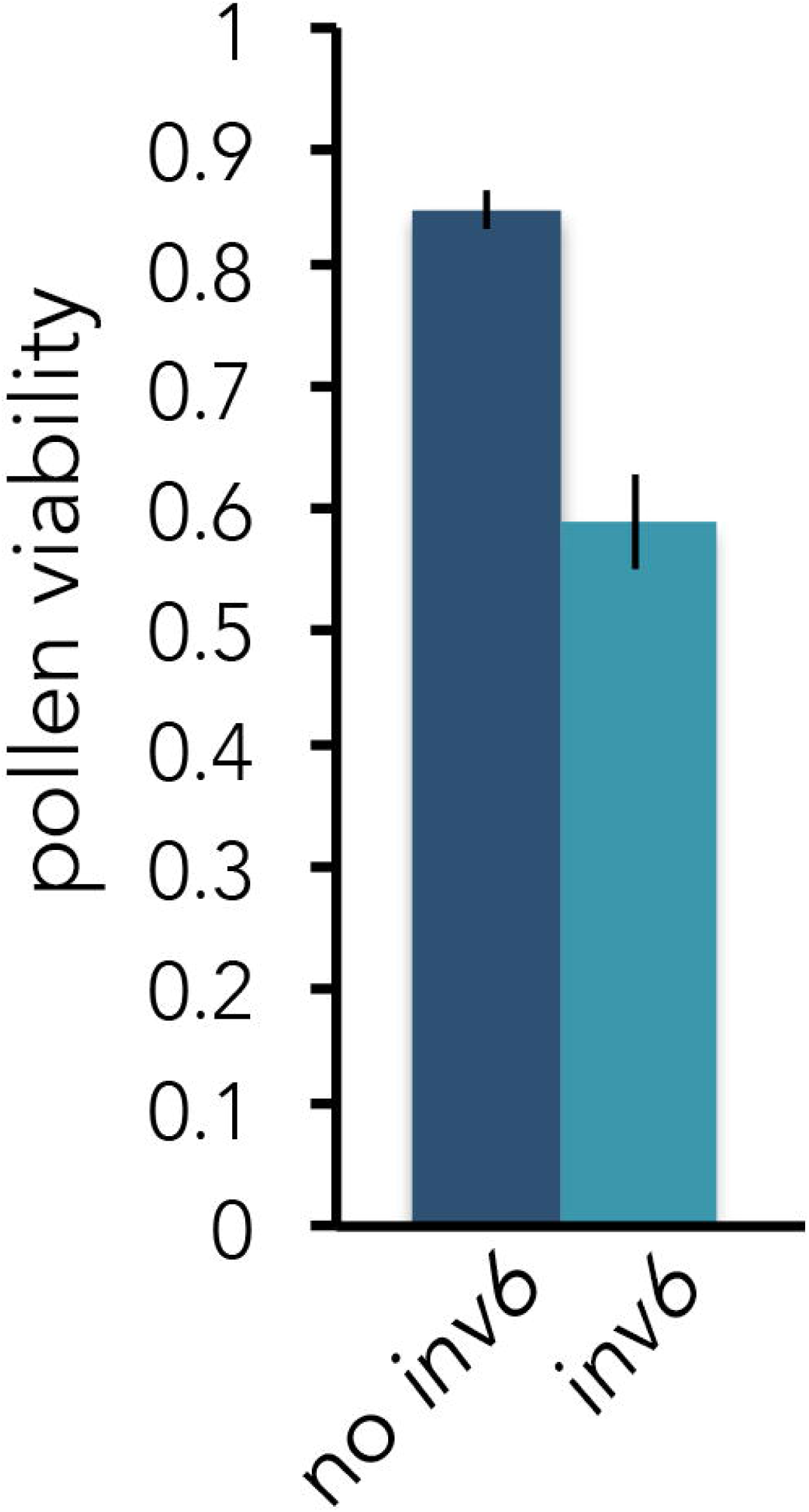
Effect of *inv6* genotype on pollen viability (mean +/− 1 SE) of wild Iron Mountain M. guttatus plants (2007; N = 177). Individuals were assigned to genotypic categories (*inv6*, no *inv6*) based on their alleles at the diagnostic markers e423 and e723. An *inv6* assignment indicates a heterozygous individual, as the two *inv6* homozygotes in the dataset were excluded.

In contrast to *inv6*’s negative effects on seedset per flower in the greenhouse and on male fertility in both environments, we detected only positive effects of *inv6* on female reproductive traits (flower, fruit, and total seed number) in the field (Figure 5). Most strikingly, for the two years in which we had flower counts (2012 and 2013), *inv6* carriers produced significantly more flowers than alternative genotypes (GLM, LR X^2^ = 7.63; *P* = 0.006, N = 889). In those two years, increased flower production translated into increased fruit set (*P* = 0.04); however, the effect on fruit number was marginally non-significant across all four years (*P* = 0.12, N = 1278). In contrast to the greenhouse experiments, we saw no *inv6* effect on seeds per fruit (*P* = 0.78, N = 1278). However, across all four years, there was a significant positive effect of *inv6* on log(total seeds) (*P* = 0.047). Although the year x *inv6* interaction was not significant for seed number (*P* = 0.13), there did seem to be a pattern. In 2010 and 2012, which had relatively low fecundity (population mean = 43 and 53 seeds, respectively), *inv6* appeared beneficial, whereas it had slightly negative effects in 2011 and 2013 (population mean = 110 and 227, respectively) (Figure 4). Grouping years by quality (2011 and 2013 good, 2010 and 2012 bad), there was a significant quality x *inv6* interaction (*P* = 0.02), as well as a significant effect of *inv6* (*P* = 0.03), in the full model.

**Figure 5.**
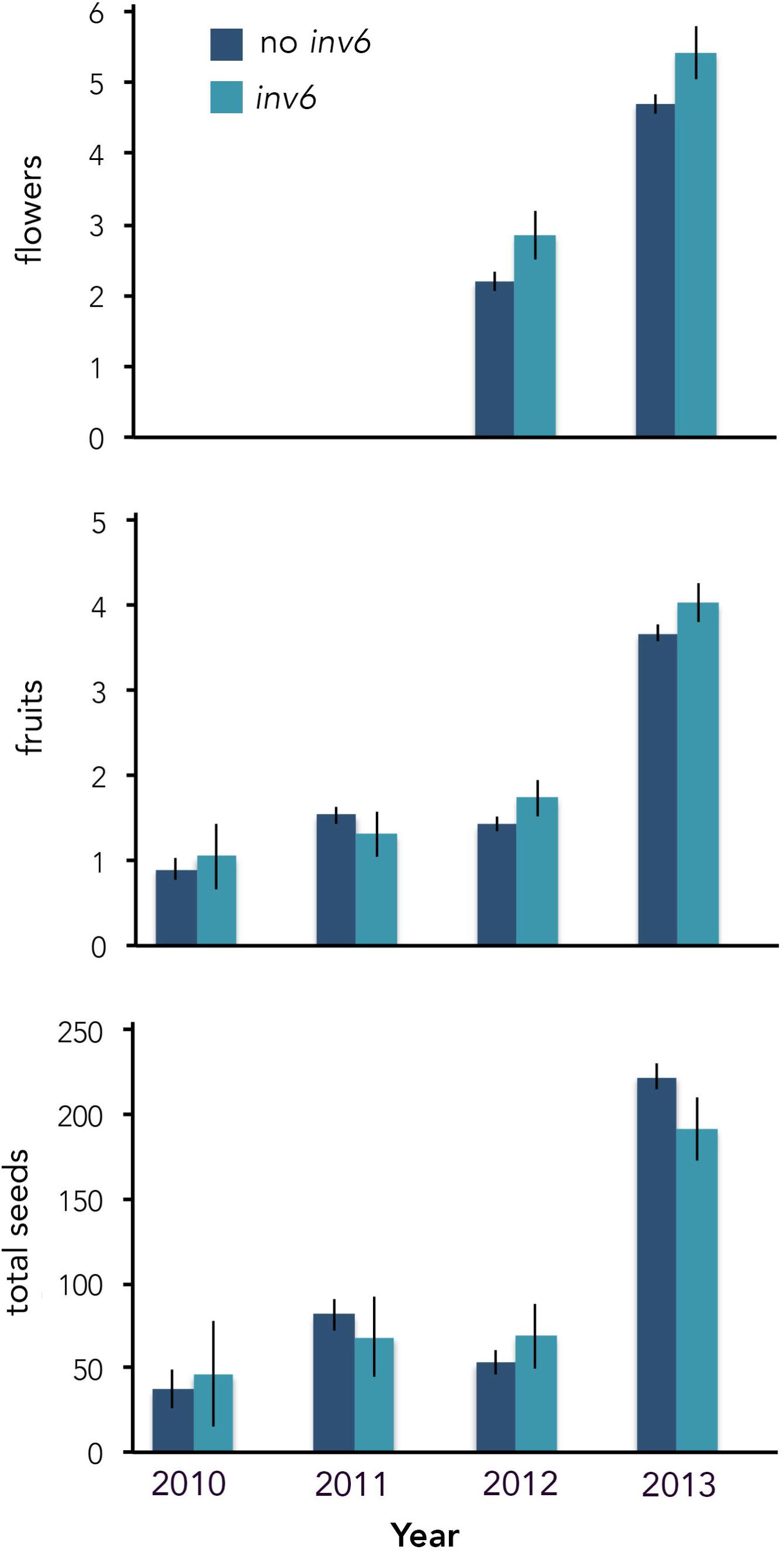
Effects of *inv6* genotype on female fitness components (mean +/− 1 SE) of wild Iron Mountain *M. guttatus* plants (N = 1248). Individuals were assigned to genotypic categories (inv6, no inv6) based on their alleles at the diagnostic marker e423. There were only a few *inv6* homozygotes in the entire four-year dataset (not enough to include in the statistical analyses), so an *inv6* assignment indicates a heterozygous individual. We show raw means and standard errors here, but the statistical tests in the text were done in a GLM framework (Poisson, log-link for fruits and flowers) or with log-transformed values (normal, identity-link for seeds).

## DISCUSSION

Understanding the genetic basis of fitness variation is key for addressing many questions in evolutionary biology, as well as applied issues in human health and agriculture. Chromosomal rearrangements such as inversions, which can bind hundreds of genes into single genetic loci with diverse effects, by preventing recombination in heterokaryotypes. They are increasingly recognized as contributors to local adaptation and speciation. Inversion polymorphisms may also be an important component of complex trait variation within populations, but segregating inversion polymorphisms are little explored beyond a few supergenes and selfish elements (Ford 1971; Thomas et al. 2008; Thompson and Jiggins 2014). Here, we genetically and phenotypically characterize a novel intermediate-frequency inversion polymorphism in the Iron Mountain population of *Mimulus guttatus* (yellow monkeyflower). We identified *inv6* on the basis of no recombination among markers on LG6 spanning over 40cM in a freely recombining cross and defined it as a distinct and apparently recently derived haplotype. We demonstrate that *inv6* has strong and partially recessive deleterious effects on male and female fertility traits in the greenhouse and negative effects on pollen fertility in wild plants, but positive effects on lifetime female fitness likely mediated through increases in flower production. These results support the hypothesis that inversion polymorphisms may commonly segregate within populations because they bind deleterious and beneficial variants into single genetic loci with balanced fitness effects.

***Why is* inv6 *polymorphic?*** The initial mapping experiment that identified *inv6* revealed only strongly negative fitness effects (Figure 2). Despite this, extensive field sampling indicates that *inv6* is surprisingly common in the largely outbred IM population. The observed frequency of 7-10% translates to about 50,000 copies of *inv6* among the ≈300,000 flowering adults in a typical year at IM. IM exhibits very high levels of nucleotide variation (ca. 2% genome wide) and there is no evidence of a population bottleneck that could have recently inflated the frequency of an unconditionally deleterious mutation (Puzey, Willis, and Kelly, unpublished manuscript). Although we do not yet know the full phenotypic effects of genes within *inv6*, the synthesis of data from this and several other experiments provide a partial explanation for this paradoxical abundance. Hardy-Weinberg proportions are immediately relevant to this explanation. In the field, there are 10-20x as many *inv6* heterozygotes as homozygotes. As a consequence, a slight advantage of *inv6* in heterozygotes may allow it to increase when rare even if it is strongly deleterious when homozygous.

The balance of evidence suggests variable, but on average, partially recessive deleterious effects for *inv6*, not the underdominance expected for sterility caused by chromosomal differences per se, i.e. gametes with duplications/deletions resulting from crossovers in inversion loops (White 1969). One of three crosses (c2) presented here revealed recessive gene action, while the other two were more nearly additive (Figure 2). This variability is not surprising, given that the alternative to *inv6* is not a single allele but many distinct haplotypes. Consistent with the recessive to additive effects we observe here, a distinct experiment involving many inbred lines from IM revealed significant partial recessivity of *inv6* in its effects on pollen number and pollen viability (see Table 2 of (Scoville et al. 2009)). Two additional experiments suggest that *inv6* is strongly deleterious in homozygotes because it is rapidly ‘purged’ when a population is experimentally inbred. Willis (1999b) initiated over 1000 independent inbred lines starting each from an outbred IM plant. After 6 generations of self-fertilization, the frequency of *inv6* was reduced to ≈ 3% (4/138, (Scoville et al. 2009)). In a distinct experiment, Bodbyl Roels and Kelly (2011) synthesized large synthetic populations by intercrossing the F_2_ populations of Figure 2 for subsequent experimental evolution. Two replicate populations were maintained at large size but compelled to self-fertilize. The initial frequency of *inv6* was 50% in each replicate, but declined to 1% and 16%, respectively, after 5 generations.

The field data for *inv6* (Figures 4-5) paint a different picture, but these estimates are based almost entirely on the heterozygous effects of the inversion (very few *inv6* homozygotes were sampled). The single year that male fitness was estimated (2007) indicates a 30% pollen viability cost in wild heterozygotes (Figure 4). This estimate is intermediate to that obtained from c3 and c4 of the replicated-F_2_ experiment (Figure 2). In contrast, field data on female fitness show that, particularly in poor years, *inv6* carriers set significantly more seeds than non-carriers. This appears to be primarily mediated through increases in flower and fruit number (i.e., through whole plant traits). A second line of evidence that heterozygous effects of *inv6* via whole-plant traits can be advantageous comes from an artificial selection experiment (Kelly 2008). Large experimental populations were founded from IM and maintained with enforced outcrossing for 10 generations. Over 10 generations, *inv6* rose from an initial frequency of 15% to ~65% in each of three independent populations, one which experienced selection for larger flowers, one for smaller flowers, as well as the unselected control (Kelly et al. 2013). The parallel increase of inv6 suggests unmeasured fitness benefits to inv6 under the conditions of that experiment. Because there was no opportunity for direct selection on flower number, no evidence of an association with flower size, and *inv6* appears costly in terms of per flower seedset in this study, we suspect that both the field and greenhouse benefits of *inv6* are mediated through other traits. One possibility is germination requirements/timing, which could both respond to inadvertent greenhouse selection, and also underlie the genotypic differences in flower number and seedset seen in the wild. Another possible advantage of *inv6* could be meiotic drive and/or pollen competition advantage. Either process would be more effective with outcrossing than selfing, thus predicting the differing outcomes from the prior studies described above. However, the replicated F_2_ mapping populations did not show transmission ratio distortion indicative of meiotic drive.

While much remains unknown about the fitness costs and benefits of *inv6*, a reasonable hypothesis is that the negative effects of *inv6* are due to partially recessive deleterious mutations captured within a novel inversion. This inversion has reached an unexpectedly high frequency owing to (perfectly) linked alleles with beneficial heterozygous effects. This interpretation is generally congruent with the model of Kirkpatrick and Barton (2006). An inversion polymorphism that has captured both advantageous and deleterious alleles is maintained if the advantageous effect of the inversion is 1) greater than its disadvantageous effects in the heterozygote and 2) smaller than the disadvantageous effects in the homozygote. Of course, these are conditions for polymorphism under almost any diploid model, inversion or not. However, inversions might greatly increase the likelihood that the conditions are met – advantageous and deleterious effects are bundled together by recombination suppression. Because the costs of *inv6* are not strictly recessive, the equilibrium frequency in the wild is predicted to be much lower than the driver *D* in the same population (Fishman and Saunders 2008; Fishman and Kelly 2015). Also, it is clear that the fitness effects and evolutionary dynamics of *inv6* depend strongly on mating system and growth conditions, and its frequency may be determined in part by fluctuating environmental conditions. In addition, we cannot yet rule out the possibility that co-adaptation among alleles within *inv6* underlie its beneficial effects, so that it is acting as a “supergene, with baggage.” Epistasis for fitness-related traits is common in *M. guttatus* (Kelly 2005; Monnahan and Kelly 2015); if positively interacting alleles are located on the same chromosome, novel inversions may often be favored because they suppress recombination among them.

*Inv6* appears recently derived and exhibits the molecular pattern of a recent partial sweep, similar to the pattern seen for the driving *D* haplotype (Fishman and Saunders 2008). *Inv6* is characterized by a specific combination of alleles at 13 markers spanning > 4.2 Mb (e25 to e480; Figure 3). This genomic region is gene-dense, containing 780 genes in the v2.0 annotation of the *Mimulus guttatus* genome. Importantly, however, *inv6* marker alleles are usually, but not always, present in non-*inv6* haplotypes as well. The latter exhibit many different combinations of alleles, indicative of free recombination in the majority of the population. Together, these observations indicate that *inv6* is the derived arrangement and that it has spread recently enough (within IM, at least) that it has accumulated little new “internal” variation via mutation. Thus, it appears that *inv6* recently captured a specific constellation of alleles, whose net positive effects in heterozygotes have carried it to substantial frequency (relative to expectation) despite substantial fitness costs.

***The genetic basis of inbreeding depression –*** Inbreeding depression, the decrease in fitness with increased homozygosity caused by mating between relatives, has been a focus of research for over 150 years (Darwin 1876; East 1908; Crow 1993). Two models have dominated thinking about inbreeding depression: dominance and overdominance. The dominance model posits that inbreeding depression is caused by recessive or partially recessive alleles with deleterious effects on fitness. Recessivity of segregating deleterious mutations is predicted because selection more rapidly eliminates additive and dominant mutations from a population. In contrast, the overdominance model states that heterozygotes have superior fitness compared to either alternative homozygote, such that fitness declines as homozygosity increases with inbreeding. At present, a preponderance of data favors mutation-selection balance of deleterious partially recessive mutations rather than overdominance to explain the bulk of inbreeding depression (Charlesworth and Willis 2009).

The major chromosomal polymorphisms that have been mapped within IM, *inv6* and the meiotic drive locus on LG11 (*D*), each generate substantial inbreeding depression. However, these polymorphisms do not conform to either the dominance or overdominance model. These loci exhibit partially-recessive deleterious effects (like the dominance model) but intermediate allele frequencies (like the overdominance model). Because of the latter feature, these polymorphisms generate considerable inbreeding depression and also genetic variance for fitness. For both *D* and *inv6*, the existence of such intermediate frequency deleterious variation depends on structural variants that prevent (at least in the short-term) recombination from breaking up the association between alleles with positive and negative effects. Otherwise, the alleles with deleterious effects (particularly if not entirely recessive) should be driven to low frequency by selection. Both deleterious recessive mutations and structural mutations are common; in combination with rare beneficial variants or driving selfish elements, they may often contribute to balanced polymorphism.

Apart from *D* and *inv6*, the data from the replicated F_2_ experiment are mainly consistent with the dominance model (inbreeding depression maintained by mutation-selection balance). We mapped over 20 addition QTLs for male fertility (Supplemental Table S2). Nearly all were mapped in only one of the three crosses. This is consistent with the prediction that deleterious alleles should be rare in the population, and as a consequence, each such allele was sampled into only one founding parent of the six used to generate the mapping populations. A few QTLs did exhibit apparent overdominance. However, these QTLs were also unique to a mapping population, contrary to expectations for alleles maintained at intermediate population frequencies by balancing selection. The level of mapping resolution in the replicated F_2_ experiment cannot distinguish true overdominance from pseudo-overdominance (deleterious alleles linked in repulsion phase). The remaining QTLs exhibit average partially recessive gene action of the low allele. The average dominance coefficient (excluding the chromosomal rearrangements) of h=0.18 is very close to the previous estimate of h~0.15 (Willis 1999a). It is likely that many additional deleterious mutations are segregating in IM, but are yet unmapped in because the individual effects are below our detection limit.

***Conclusion—***Modern tools of genetic mapping, combined with population sampling, provide a straightforward and productive means to investigate the maintenance of genetic variation in fitness. This work contributes to a larger effort to identify genetic components of fitness variation within the Iron Mountain population of *M. guttatus* and reveal their individual (and apparently complex) histories. Balancing selection facilitated by recombination suppression in inversions may be an unexpectedly significant and general factor in the maintenance of fitness variation within populations, in keeping with the prominent role of inversions in speciation and divergence.

## Acknowledgements

We thank Arpiar Saunders, Tyler Huggins, Angela Stathos, Dan Crowser, Becky Fletcher, Katie Zarn, and Mariah McIntosh for assistance with field collections, plus counting of pollen and seeds and genotyping of markers. The research was supported by grant NIH GM073990 to JK and JW and by NSF DEB-0918902 to LF.

## Cited

Balanyà, J., L. Serra, G. W. Gilchrist, and R. B. Huey. 2003. Evolutionary pace of chromosomal polymorphism in colonizing populations of Drosophila subobscura: an evolutionary time series. Evolution 57: 1837–1845.

Bodbyl Roels, S. A. and J. K. Kelly. 2011. Rapid evolution caused by pollinator loss in Mimulus guttatus. Evolution 65: 2541–2552.

Burt, A. and R. Trivers. 2008. Genes in Conflict. Belknap Press.

Charlesworth, D. and J. H. Willis. 2009. The genetics of inbreeding depression. Nat Rev Genet 10: 783–796.

Cheng, C., B. J. White, C. Kamdem, K. Mockaitis, C. Costantini, M. W. Hahn, and N. J. Besansky. 2012. Ecological Genomics of Anopheles gambiae Along a Latitudinal Cline: A Population-Resequencing Approach. Genetics 190: 1417–1432.

Coluzzi, M., A. Sabatini, A. della Torre, M. A. Di Deco, and V. Petrarca. 2002. A Polytene Chromosome Analysis of the Anopheles gambiae Species Complex. Science 298: 1415–1418.

Crow, J. F. 1993. Mutation, mean fitness, and genetic load. Oxford surveys in evolutionary biology 9: 3–42.

Darwin, C. R. 1876. The effects of cross- and self-fertilisation in the vegetable kingdom. John Murray, London.

Dobzhansky, T. 1970. Genetics of the evolutionary process. Columbia University Press, New York.

East, E. M. 1908. Inbreeding in corn. Rep. Conn. Agric. Exp. Sta.:419–428.

Fang, Z., T. Pyhäjärvi, A. L. Weber, R. Dawe, J. C. Glaubitz, J. D. J. S. Gonzalez, C. Ross-Ibarra, J. F. Doebley, P. L. Morrell, and J. Ross-Ibarra. 2012. Megabase-Scale Inversion Polymorphism in the Wild Ancestor of Maize. Genetics 191: 883–894.

Feder, J. L., J. B. Roethele, K. Filchak, J. Niedbalski, and J. Romero-Severson. 2003. Evidence for Inversion Polymorphism Related to Sympatric Host Race Formation in the Apple Maggot Fly, Rhagoletis pomonella. Genetics 163: 939–953.

Fishman, L., A. J. Kelly, E. Morgan, and J. H. Willis. 2001. A genetic map in the Mimulus guttatus species complex reveals transmission ratio distortion due to heterospecific interactions. Genetics 159: 1701–1716.

Fishman, L. and J. K. Kelly. 2015. Centromere-associated meiotic drive and female fitness variation in Mimulus. Evolution 69: 1208–1218.

Fishman, L. and A. Saunders. 2008. Centromere-Associated Female Meiotic Drive Entails Male Fitness Costs in Monkeyflowers. Science 322: 1559–1562.

Fishman, L., A. Stathos, P. Beardsley, C. F. Williams, and J. P. Hill. 2013. Chromosomal rearrangements and the genetics of reproductive barriers in mimulus (monkey flowers). Evolution 67: 2547–2560.

Fishman, L., J. H. Willis, C. A. Wu, and Y. W. Lee. 2014. Comparative linkage maps suggest that fission, not polyploidy, underlies near-doubling of chromosome number within monkeyflowers (Mimulus; Phrymaceae). Heredity 112: 562–568.

Ford, E. B. 1971. Ecological genetics. Chapman and Hall, London.

Gilburn, A. S., and T. H. Day. 1999. Female mating behaviour, sexual selection and chromosome I inversion karyotype in the seaweed fly, Coelopa frigida. Heredity 82: 276–281.

Hermann, K., U. Klahre, M. Moser, H. Sheehan, T. Mandel, and C. Kuhlemeier. 2013 Tight genetic linkage of prezygotic barrier loci creates a multifunctional speciation island in Petunia. Curr. Biol. 23: 873–877.

Hoffmann, A. A., and L. H. Rieseberg. 2008. Revisiting the Impact of Inversions in Evolution: From Population Genetic Markers to Drivers of Adaptive Shifts and Speciation? 39: 21–42.

Jones, F. C., M. G. Grabherr, Y. F. Chan, P. Russell, E. Mauceli, J. Johnson, R. Swofford, M. Pirun, M. C. Zody, S. White, E. Birney, S. Searle, J. Schmutz, J. Grimwood, M. C. Dickson, R. M. Myers, C. T. Miller, B. R. Summers, A. K. Knecht, S. D. Brady, H. Zhang, A. A. Pollen, T. Howes, C. Amemiya, E. S. Lander, F. Di Palma, K. Lindblad-Toh, and D. M. Kingsley. 2012. The genomic basis of adaptive evolution in threespine sticklebacks. Nature 484: 55–61.

Kelly, A. J., and J. H. Willis. 1998. Polymorphic microsatellite loci in Mimulus guttatus and related species. Mol. Ecol. 7: 769–774.

Kelly, J. K. 2005. Epistasis in Monkeyflowers. Genetics 171: 1917–1931.

Kelly, J. K. 2008. Testing the rare alleles model of quantitative variation by artificial selection. Genetica 132: 187–198.

Kelly, J. K., B. Koseva, and J. P. Mojica. 2013. The Genomic Signal of Partial Sweeps in Mimulus guttatus. Genome Biology and Evolution 5: 1457–1469.

Kelly, J. K., A. Rasch, and S. Kalisz. 2002. A method to estimate pollen viability from pollen size variation. American Journal of Botany 89: 1021–1023.

Kelly, J. K., and J. H. Willis. 2002. A manipulative experiment to estimate bi-parental inbreeding in Monkeyflowers. International Journal of Plant Science 163: 575–579.

Kondrashov, A. S. 1988. Deleterious Mutations and the Evolution Of Sexual Reproduction. Nature 336: 435–440.

Krimbas, C. B., and J. R. Powell. 1992 Drosophila Inversion Polymorphism. CRC Press, Boca Raton, FL.

Lowry, D. B., and J. H. Willis. 2010. A Widespread Chromosomal Inversion Polymorphism Contributes to a Major Life-History Transition, Local Adaptation, and Reproductive Isolation. PLoS Biol 8:e1000500.

Monnahan, P. J., and J. K. Kelly. 2015. Naturally segregating loci exhibit epistasis for fitness.

Noor, M. A. F., K. L. Grams, L. A. Bertucci, and J. Reiland. 2001. Chromosomal inversions and the reproductive isolation of species. Proc. Natl. Acad. Sci. USA 98: 12084–12088.

Rieseberg, L. H., J. Whitton, and K. Gardner. 1999. Hybrid zones and the genetic architecture of a barrier to gene flow between two sunflower species. Genetics 152: 713–727.

Schwander, T., R. Libbrecht, and L. Keller. 2014. Supergenes and Complex Phenotypes. Current Biology 24:R288–R294.

Scoville, A., Y. W. Lee, J. H. Willis, and J. K. Kelly. 2009. Contribution of chromosomal polymorphisms to the G-matrix of Mimulus guttatus. New Phytologist 183 803–815.

Stam, P. 1993. Construction of integrated genetic linkage maps by means of a new computer package: Join Map. Plant J 3: 739–744.

Sweigart, A., K. Karoly, A. Jones, and J. H. Willis. 1999. The distribution of individual inbreeding coefficients and pairwise relatedness in a population of Mimulus guttatus. Heredity 83: 625–632.

Thomas, J. W., M. Cáceres, J. J. Lowman, C. B. Morehouse, M. E. Short, E. L. Baldwin, D. L. Maney, and C. L. Martin. 2008. The chromosomal polymorphism linked to variation in social behavior in the white-throated sparrow (Zonotrichia albicollis) is a complex rearrangement and suppressor of recombination. Genetics 179: 1455–1468.

Thompson, M. J., and C. D. Jiggins. 2014. Supergenes and their role in evolution. Heredity 113: 1–8.

Vickery, R. K. 1978. Case studies in the evolution of species complexes in Mimulus. Evol Biol 11: 405–507.

White, M. J. D. 1969. Chromosomal rearrangements and speciation in animals. Annu. Rev. Genet. 3: 75–98.

Willis, J. H. 1993. Partial self fertilization and inbreeding depression in two populations of Mimulus guttatus. Heredity 71: 145–154.

Willis, J. H. 1999a. Inbreeding load, average dominance, and the mutation rate for mildly deleterious alleles in Mimulus guttatus. Genetics 153: 1885–1898.

Willis, J. H. 1999b. The role of genes of large effect on inbreeding depression in Mimulus guttatus. Evolution 53: 1678–1691.

Wu, C. A., D. B. Lowry, A. M. Cooley, K. M. Wright, Y. W. Lee, and J. H. Willis. 2007. Mimulus is an emerging model system for the integration of ecological and genomic studies. Heredity 100: 220–230.

Yandell, B. S., T. Mehta, S. Banerjee, D. Shriner, R. Venkataraman, J. Y. Moon, W. W. Neely, H. Wu, R. von Smith, and N. Yi. 2007. R/qtlbim: QTL with Bayesian Interval Mapping in experimental crosses. Bioinformatics 23: 641–643.

Yi, N., S. Banerjee, D. Pomp, and B. S. Yandell. 2007. Bayesian Mapping of Genomewide Interacting Quantitative Trait Loci for Ordinal Traits. Genetics 176: 1855–1864.

